# Colour morphs as alternative solutions to the trade-off predicted by the Immuno-Competence Handicap Hypothesis

**DOI:** 10.1101/2024.11.26.625393

**Authors:** Roberto Sacchi, Alan J. Coladonato, Stefano Scali, Marco A. L. Zuffi, Rupert Palme, Marco Mangiacotti

## Abstract

Colour morphs in polymorphic species represent a suite of heritable traits controlled by separate sets of loci, which correspond to alternative peaks in the fitness surfaces. Pleiotropic effects of hormones have been hypothesized to be involved in the maintenance of morphs, but no experiments have been done to test it so far. In this study, we experimentally increased testosterone plasma levels in colour morph of the common wall lizards to test if white and yellow morphs follow two alternative strategies imposed by trade-off depicted in the Immuno-Competence Handicap Hypotheses. Yellow males invest in a more aggressive behaviour at the expense of a reduced immune function, whereas white males invest in greater survival at the expense of a reduced competitiveness. Plasma T levels were increased through transdermal administration. Before and after treatment, we measured i) the immune-response of lizards through phytohaemoagglutinin (PHA), and ii) the aggressive response by the introduction of a small mirror into the plastic enclosure to mimic the intrusion of a stranger male in its own territory. The hormone administration caused a higher immune-suppression in white than yellow males, and a switch to less aggressive behaviour against the mirrored image. Overall, increased plasma testosterone levels resulted in the disappearance of differences between morphs of both immune-response and aggressive behaviour. These results supplied experimental evidence of the existence of morph-specific strategies in common wall lizards depending on the investment in territorial aggression or in a longer survival, as alternative solutions to the trade-off predicted by the Immuno- Competence Handicap Hypothesis.

## INTRODUCTION

Understanding the origin of colour polymorphism, i.e. the co-occurrence within the same population of genetically inherited colour variants (morphs, *1*, *2*), is a major challenge in evolutionary ecology. Morphs are traditionally interpreted as alternative peaks in the fitness surfaces promoted by disruptive selection in response to specific life history trade-offs (Sinervo and Lively 1996; Svensson et al. 2001; McKinnon and Pierotti 2010). Therefore, morphs represent a suite of heritable traits controlled by separate sets of loci, including those involved in the expression of the alternative colourations, which affect fitness in an interactive way (Whitlock et al. 1995; Sinervo and Svensson 2002). These favourable genetic correlations are assembled by correlative selection, which promotes linkage disequilibrium among the loci controlling the expression of the optimal combinations of the phenotypic traits that are identifiable as morphs (Sinervo and Svensson 2002). There is general agreement that correlational selection is the first step in building-up the adaptive genetic associations (Svensson 2017), but other selective forces, such as frequency-dependent selection, are needed for maintaining alleles combinations within morphs once assembled (Sinervo and Lively 1996; Sinervo and Svensson 2002).

Pleiotropic effects of the colour locus have been also hypothesized to be involved in the maintenance of morphs, for example, hormones which can affect multiple traits including colour, behaviours, physiology and reproductive traits (Sinervo et al. 2000; Comendant et al. 2003). Indeed, hormones trigger a set of behavioural responses which make individuals able to deal with environmental stressors (e.g. glucocorticoids) or sexual competition (e.g. testosterone). On the other hand, when hormone levels remain high for long time, negative consequences can occur including reduced immune-competence (Munck et al. 1984; Kurtz et al. 2007). Therefore, a trade-off between immune-response and other functions relevant for individual survival and reproduction can be generated, ultimately affecting the individual fitness (Folstad and Karter 1992). Since individuals show considerable variability on how they respond to stressors, and how behaviour and physiology adjust to these stressors, frequency dependent selection could promote interactions between resource allocation in response to the trade-off and morphs (Comendant et al. 2003).

Here we tested these hypotheses on the colour morphs of the common wall lizard (*Podarcis muralis*) through experimental manipulations of plasma testosterone (T) levels. This species is a small diurnal lizard widespread in central and southern Europe (Sillero et al. 2014), which exhibits three discrete colour morphs (yellow, white and red) genetically determined in both sexes (Sacchi, Scali, et al. 2007; Andrade et al. 2019). In the last decades morph-specific patterns in several life-history traits were found in this species (Calsbeek et al. 2010; Galeotti et al. 2013; Pérez i de Lanuza et al. 2013; Scali et al. 2013; Sacchi et al. 2015; Scali et al. 2015; Mangiacotti et al. 2019). In particular, yellow males are immunosuppressed (Sacchi, Rubolini, et al. 2007; Sacchi, Mangiacotti, et al. 2017) compared to the other morphs, and bear high plasma T levels at the beginning of breeding season displaying a stronger subsequent decline towards the end (Sacchi, Scali, et al. 2017). The seasonal trends of the hormone match with the seasonal pattern of the aggressive behaviour (Coladonato et al. 2020). In fact, the absolute T levels did not actually differ among morphs, but each morph reaches the hormone peak in different periods of the season, and the peak in the aggressive response follows nearly a month later.

Consequently, yellow males show higher T level-correlated aggressiveness than white males at the beginning of the breeding season, whereas the opposite occurs in the second part of the season, when white males become more aggressive (Sacchi, Scali, et al. 2017; Coladonato et al. 2020).

Based on all above data, we hypothesized that white and yellow morphs follow two distinct, alternative strategies as two opposite solutions for the constraints imposed by the trade-off depicted in the Immuno-Competence Handicap Hypotheses (ICHH, 12). High testosterone plasma level can decrease immune functions, favours parasite infections, stimulates risky behaviours, and thus diminishes survival (Olsson et al. 2000; Klukowski and Nelson 2001; Cox and John-Alder 2007), and there is therefore no way for males of maximizing stamina and aggressive behaviors at once. Accordingly, yellow males play a risky strategy, where males invest in more aggressive behavior at the expense of a reduced immune function, whereas a conservative strategy is played by not-yellow males (i.e. white and red ones), where the fitness is achieved with greater survival.

The effect of experimentally T levels manipulation is well documented in many lizard species: treated males obtain larger and higher-quality home ranges, and are more successful in fighting against opponents (Marler and Moore 1989; DeNardo and Sinervo 1994). On the other hand, increasing plasma T levels leads to a decrease of cell-mediated immunity (Olsson et al. 2000; Oppliger et al. 2004). The manipulation of T plasma level has been successfully tested in *Podarcis* lizards (Oppliger et al. 2004; Baeckens and Van Damme 2018), but no previous work has looked for morph-specific patterns of response to T manipulation. In the present study, we manipulated plasma T levels in yellow and white males of the common wall lizard in order to experimentally test the interplay between the trade-off predicted by ICHH and colour morphs. If the risky and conservative strategies are followed by yellow and white males respectively, we expect to observe morph-specific pattern in both immune-competence, and aggressive response. Notably, if white males preferentially allocate most resources in survival, we expect to observe a more intense immune-suppression in white than in yellow males. On the other hand, if immune function affects aggressive behaviour, we expect to observe a depression of aggressive response, more pronounced in white than yellow males.

## METHODS

### Lizard collection and housing

Forty adult males of common wall lizards were captured during March 2019 (20^th^ – 26^th^) in and around the town of Pavia (Northern Italy, 45°11′N, 9°9′E). Only pure white (n=20) and yellow (n=20) morphs were collected, according to Sacchi et al. (Sacchi et al. 2013). The SVL of each individual was measured to the nearest 0.5 mm and weighed to the nearest 0.1 g. We individually housed lizards in 20 × 30 × 20 cm transparent plastic boxes, with a brick as shelter/basking site, a small bowl with water, a UVB 5% lamp turned on 8 hours/day (Sylvania F30W Reptistar T8 UVB 5%), and heating pads as hot spot. We provided mealworms as food (one mealworm/day), which were dusted with vitamin and calcium supplements two times for week. A minimum acclimation period of one week was given before starting trials, and we released all lizards at their capture sites following experimental trials. No lizard was injured or killed during the study, and all lizards looked healthy at release.

### Manipulation of testosterone

Plasma T levels were manipulated according to the non-invasive technique already proposed for lacertid species (Knapp and Moore 1997; Belliure et al. 2004; Oppliger et al. 2004), and successfully used in *P. muralis* (Baeckens et al. 2016).

Following this last study, we increased plasma T levels through transdermal administration of a mixture of the steroid hormone and sesame oil to the lizard’s dorsal skin. Lipophilic molecules such as testosterone are able to pass through lizard scales into blood stream (Baeckens et al. 2016). The administered T- dose was the same as in Baeckens et al. (Baeckens et al. 2016), obtained by diluting testosterone (4- androsten-17b-ol-3-one; Sigma #86500) in commercial sesame to a concentration of 4 µg/µL. Males received 4 µL of the hormone dilution every two days over four consecutive weeks. We applied a droplet of the hormone solution on the back of lizards early in the morning, before turning on the UVB lamp and heating pad, when individuals were not in activity, to minimize the stress. To ensure that the treatment successfully increased plasma T levels, we used a non-invasive steroid analysis based on faecal dropping (Palme 2005; Palme et al. 2013). We collected faecal drops for each individual before the hormonal treatment and at the end of the experiment.

Samples were frozen and stored in individual labelled Eppendorf tubes (0.5 mL) at -20°C. Concentrations of faecal testosterone metabolites (TMs) were measured with a testosterone enzyme immunoassay (EIA), previously utilized in lizards droppings (Baeckens et al. 2016). For details of the EIA see Auer et al. (Auer et al. 2020).

### Behavioural experiments

Each individual performed double behavioural experiments before and after the hormone treatment. We measured the aggressive response of a focal male by the introduction of a small mirror (15 × 15 cm) into the plastic enclosure to mimic the intrusion of a stranger male in its own territory. We had previously shown that common wall lizards perceive their own mirror image as a rival, and behave aggressively in response (Scali et al. 2019; Coladonato et al. 2020; Sacchi et al. 2021). This procedure allows the experimenters to control for the effects of differences in size and motivation between opponents on the aggressive response of the focal male since the mirror image exhibits the same size and motivation of the male (Sacchi et al. 2021). Before starting a trial, we removed the water bowl, and put a partition dividing the arena into two halves. We then placed the mirror at the far end of the half without the lizard. After 5 minutes we assumed focal lizards has habituated to disturbance and removed the partition, thereby allowing the lizard to interact with the mirror. To avoid visual disturbance during the trials, the four sides of the arena were externally covered by opaque, white plastic panels. Before each trial, the male was heated for two minutes using a 75 W halogen infra-red lamp (Reptiles-Planet.com) positioned 40 cm above the arena. After switching off the lamp, the mean (± SD) body temperature of males just before starting the trial (measured with a handheld infra-red thermometer Lafayette TRP-39, Lafayette Instrument Co., Lafayette, Indiana, USA; sensitivity: 0.1 °C; precision: ± 2%) was 37.6 ± 1.8 °C. The lizard movements were recorded using a webcam (Microsoft LifeCam HD 3000) mounted on an easel, 60 cm above the arena, and connected to a laptop by a 3 m cable. Recording was managed by Free2X software v1.0.0.1 (freely available at: http://www.free2x.com/webcam-recorder/), setting quality to 800 × 600 pixels and 15 frames per second (fps). Recording duration was set to 15 minutes, following the first lizard movement (i.e., tongue flicking, head movement, etc.). Room temperature was set to 28 °C to reduce thermal loss during the trials.

Trials took place between 10 a.m. and 2 p.m., and the order of morphs was randomized to control for potential effects of day-time. We repeated a trial the subsequent day if the lizard did not move after 10 min from the start. The first series of trials was performed between 3^rd^ and 18^th^ April and the second series between 8^th^ and 23^th^ June.

### Response variables

We used BORIS (Friard and Gamba 2016) to analyse videos and extract four response variables. The first three variables were used to assess the aggressive behaviour responses (Coladonato et al. 2020; Ficetola et al. 2021): (i) the number of times that lizards entered in the half of the enclosure containing the mirror (Nmirror), (ii) the time (seconds) spent in the half of the enclosure with the mirror (Time), and (iii) the total number of bites against the mirrored image (Bites). These three variables can be interpreted as increasing levels in a rank of aggression (see Sacchi et al. (Sacchi et al. 2021) for details). The fourth variable was the tongue flicks rate (number of flicks divided by Time) measured in the half of the enclosure containing the mirror (TF). This variable evaluated the basal explorative behaviour of each individual when facing a potential contestant. For simplicity, we hereafter refer to Bites, Time and Nmirror as forms of “aggressive behaviour”, and to TF as “exploratory behaviour”. All response variables achieved normality (Bites required a log-transformation), and were weakly correlated with each other (Spearman correlation coefficient: |*rs*|<0.52).

### Immune-response test

*In vitro* activation of lymphocytes enabled us to repeatedly challenge the immune system of the same individual at different temperatures preventing the adaptive immune response to form an immunological memory (Sacchi, Scali, et al. 2017). We used it to evaluate the change in the immune function of each individual in response to the hormonal treatment (i.e., before and after the treatment). Blood samples (20µL) were collected in heparinized capillary tubes from the post-orbital sinus (MacLean et al. 1973), and inoculated it in 15ml of RPMI 1640 medium supplemented with 10% bovine serum. Then, we divided the cell suspension into two 7ml sub-cultures, one of which was inoculated with 1% PHA solution (PHA-P Sigma, 50mg in 10ml of phosphate-buffered saline) (Oppliger et al. 2004; Sacchi, Scali, et al. 2017). The last 1ml of solution was used to assess starting lymphocyte concentration using a Neubauer chamber. Each sub-culture was then distributed in two 1.5 ml culture tubes, and incubated at 32°C for three days. Afterward, cells were collected, re-suspended and newly counted. This second count involved only proliferating lymphocytes. Colony-forming units (CFU) and the total T-cells per ml to the corresponding were assessed and stimulation was evaluated by the fold-change of the PHA sample with respect to the control (Sacchi, Mangiacotti, et al. 2017).

### Statistical analysis

First at all, we verified whether the treatment has actually increased the faecal T concentration using a random intercept linear mixed-effects model (LMM), in which the fixed effects were the treatment (pre and post), the morph and their interaction. The individual entered the model as random effect.

Secondly, to examine if the aggressive response of lizards changed according to the hormonal treatment, we still used LMMs, one for each response variable. Fixed effects were the hormonal treatment, the morph, and their interaction to account for possible differential effect of treatment due to colour morphs. We also added SVL (after standardization to zero-mean and unite variance) as fixed effect to control for possible confounding effect due to individual size. The individuals entered the model as random effect. Bites showed a Poisson-like distribution with over dispersion (sd/mean = 29), and zero inflation, while the other variables assumed a normal distribution (One-sample Kolmogorov- Smirnov test, all P values larger than 0.05). Thus, we ran a Zero-Inflate Negative Binomial Regression for Bites, and linear mixed-effects model via Satterthwaite’s degrees of freedom for the others variable.

Thirdly, we used the same approach to search for morph-specific changes in the immune-response according to the hormonal treatment. Lymphocytes and CFU were the dependent variables in two separated LMMs, in which the fixed and the random components were the same as in the analyses of the aggressive response. Both dependent variables followed a normal distribution, and were not transformed.

Finally, we tested for the effects of the hormone treatment on the relationship between aggressive behaviour and immune-responsiveness through random intercept LMMs where Bites, Time, Nmirror and TF were the dependent variable, each in a separate model, and the three-way interaction morph×treatment×immunity was the fixed effect. Immunity was, in turn, represented by the total lymphocytes or CFU counts. Therefore, eight models were prepared, wiht individual ID as random effect on the intercept.

LMMs were fit in a Bayesian analytical framework available in the package JAGS 4.3.0 (http://mcmc-jags.sourceforge.net/), using flat priors for coefficients and intercept (μ = 0 and σ = 0.001), and uninformative half-Cauchy priors (x0 = 0, γ = 25) for both σ²error and σ²individual. For all models, Markov Chain Monte Carlo parameters were set as follows: number of independent chains = three; number of iterations = 34,000; burning = 4000; thinning = three. We checked convergence through trace plot and autocorrelation along chains and results from the posterior distribution are reported as the half sample mode (HSM, (Bickel and Frühwirth 2006) with 95% and 50% highest density intervals (HDI95, (Kruschke 2010). All analyses were done in R 3.6.1 (R Core Team 2022) using the packages R2jags (Su and Yajima 2015), modest (Poncet 2012), and HDInterval (Meredith and Kruschke 2018).

## RESULTS

One white male was excluded from the experiments because it still had not eaten after one week of acclimation, and was therefore released. Faecal samples from five lizards (1 white and 4 yellow individuals) were not heavy enough for steroid analysis, and were discarded. Consequently, the final sample included 34 individuals (18 white and 16 yellow).

### Hormonal treatment

Following the four treatment weeks, faecal testosterone metabolites (TM) concentrations doubled compared to the initial values in all males of both morphs (for both morphs: Ppost-pre>0 = 0.999, Tab. 1). No difference between morphs were observed before (Pwhite>yellow = 0.305), whereas after treatment yellow males had higher faecal TM concentration (Pwhite>yellow = 0.011, Tab. 1).

**Table 1.**
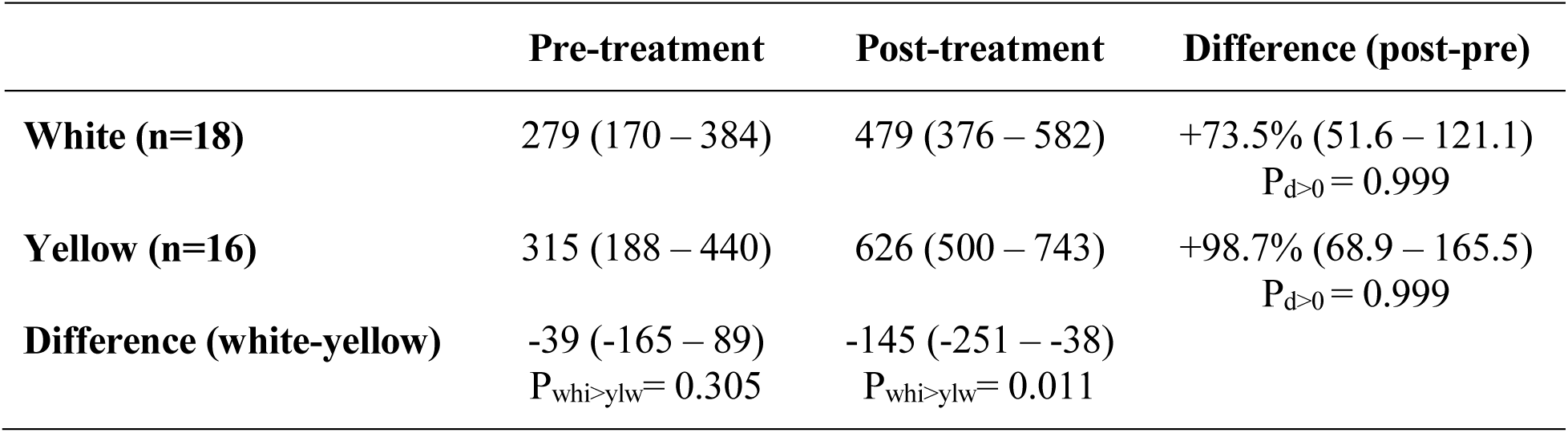
Posterior distributions of testosterone metabolite concentration (ng/g) in faecal pellets of white and yellow morphs before and after the hormonal treatment (see methods for details). HSM and HDI95 estimates (between brackets) are shown.

### Behavioural response and hormonal treatment

In 65 out of 68 trials (95.5%) lizards approached the mirror, and in 33 of them (50.8%) bit the reflected image. Focal lizards entered the half portion of the cage hosting the mirror (Nmirror) on average 7.3 ± 4.7 (range 0-19) times, whereas the time spent in the half-mirrored cage (Time) ranged from 0 to 866 s, being on average 417 ± 274 s. The mean number of tongue flicks (TF) was 43 ± 40 (0 – 196).

Following the hormonal treatment, both white and yellow morphs increased the time spent in the half-mirrored cage (Fig. 1), but white males responded more than yellow ones (relative increase, white morph: +77.4%, yellow morph: +41.8%; Tab. 2). Consequently, the difference in favour of yellow males observed before the hormonal treatment sensibly decreased in the post-treatment, when white and yellow morphs showed very similar values (Tab. 2, Fig. 1). The number of bites followed the opposite pattern (Fig. 1), and in both morphs bites decreased sensibly after the hormonal treatment (Tab. 2). Further, the extent of the decrease was similar in both morphs (white morph: -35.2%; yellow morph: -39.6%), so white males bit the mirrored image more than yellow ones irrespective of the hormone concentration. Nmirror did not appreciably varied between the two measurements within morph (Fig. 1), and the same occurred for the comparison between morphs in each measurement (Tab. 2). The TF also did not appreciably change following the hormonal treatment (Tab. 2, Fig. 1).

**Figure 1.**
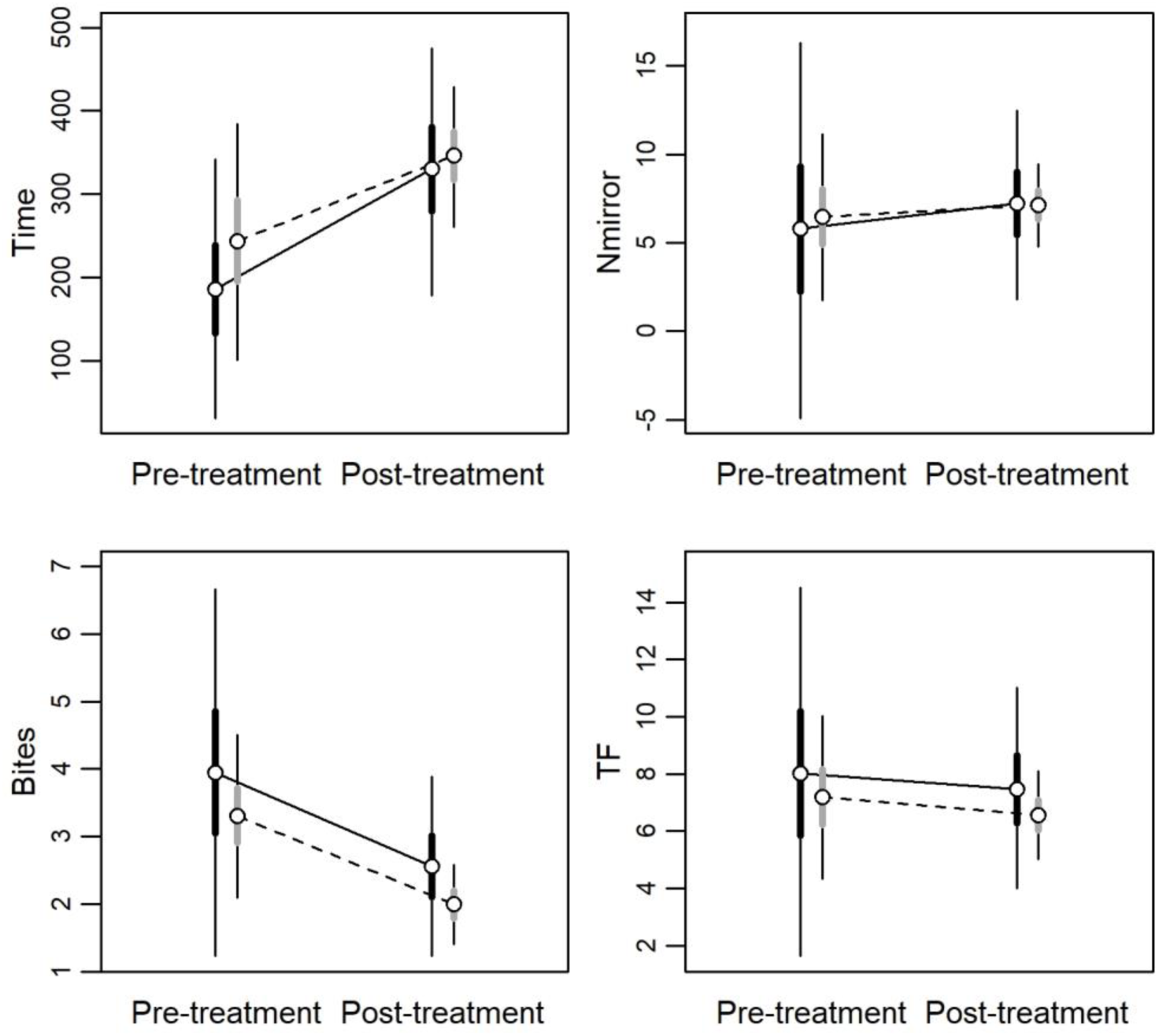
Bayesian model predictions for the aggressive and exploratory responses of common wall lizards before (pre-treatment) and following (post-treatment) the testosterone supplementation according to morph (*black and gray bars* are for white and yellow morphs, respectively). *Time*: time in seconds spent in the half enclosure containing the mirror; *Nmirror*: the number of times the lizard entered the half of the arena containing the mirror; *Bites*: the total number of bites against the mirror image (log-transformed, see text for details); *TF*: the number of tongue flicks in the half of the arena containing the mirror. Circles indicate HSM, and thick and thin lines represent HDI50 and HDI95, respectively.

**Table 2.** Posterior distributions for the aggressive and exploratory responses of common wall lizards before (pre-treatment) and following (post-treatment) the testosterone supplementation according to morph. HSM and HDI95 estimates (between brackets) are shown.

| Variable |  | Pre-treatment | Post-treatment | Difference (post-pre) |
| --- | --- | --- | --- | --- |
| Time | White | 186 (55 – 316) | 330 (204 – 451) | +144 (40 – 247)<br>$P_{d>0} = 0.998$ |
| | Yellow | 244 (124 – 363) | 346 (275 – 416) | +102 (23 – 181)<br>$P_{d>0} = 0.984$ |
| | Difference | 58 (-45 – 161)<br>$P_{ylw>whi} = 0.825$ | 16 (-65 – 101)<br>$P_{ylw>whi} = 0.622$ | |
| Nmirror | White | 5.82 (-3.09 – 14.4) | 7.22 (2.73 – 11.6) | +1.41 (-3.91 – 6.78)<br>$P_{d>0} = 0.670$ |
| | Yellow | 6.46 (2.53 – 10.4) | 7.12 (5.15 – 9.06) | +0.67 (-2.90 – 2.81)<br>$P_{d>0} = 0.685$ |
| | Difference | 0.64 (-4.94 – 6.34)<br>$P_{ylw>whi} = 0.578$ | -0.10 (-2.90 – 2.81)<br>$P_{ylw>whi} = 0.481$ | |
| Bites | White | 3.95 (1.70 – 6.19) | 2.56 (1.45 – 3.67) | -1.39 (-2.77 – -0.02)<br>$P_{d>0} = 0.048$ |
| | Yellow | 3.31 (2.30 – 4.29) | 2.00 (1.50 – 2.48) | -1.31 (-1.92 – -0.69)<br>$P_{d>0} < 0.001$ |
| | Difference | -0.64 (-2.08 – 0.83)<br>$P_{ylw>whi} = 0.229$ | -0.57 (-1.28 – 0.15)<br>$P_{ylw>whi} = 0.093$ | |
| TF | White | 8.02 (2.65 – 13.4) | 7.48 (4.58 – 10.4) | -0.55 (-3.72 – 2.66)<br>$P_{d>0} = 0.385$ |
| | Yellow | 7.19 (4.82 – 9.55) | 6.55 (5.27 – 7.84) | -0.63 (-2.05 – 0.79)<br>$P_{d>0} = 0.228$ |
| | Difference | -0.85 (-4.28 – 2.63)<br>$P_{ylw>whi} = 0.343$ | -0.91 (-2.81 – 0.96)<br>$P_{ylw>whi} = 0.207$ | |

### Immune-response and hormonal treatment

The phytohemagglutinin (PHA) injection actually stimulated the lizards’ immune system in pre- and post-hormonal treatment, as the fold-changes (Tab. 3) were systematically higher than one in all cases for both the total lymphocyte count and the colony-forming units (CFU; Pfold-change>1 > 0.959). According to lymphocytes values, white males showed higher immune-competence than yellow ones, both before and after the hormonal treatment (Tab. 3), even if the difference was reduced by two thirds following the testosterone administration (Tab. 3, Fig. 2). This reduction in the difference between morphs was entirely due to white males, which showed an evident immunosuppression due to the hormonal treatment (Fig. 2). Compared to the initial values, the immune-response of white males decreased by about 13% after testosterone administration (Tab. 3), whereas the immune-response of yellow males showed no change at all (Tab. 3).

**Figure 2.**
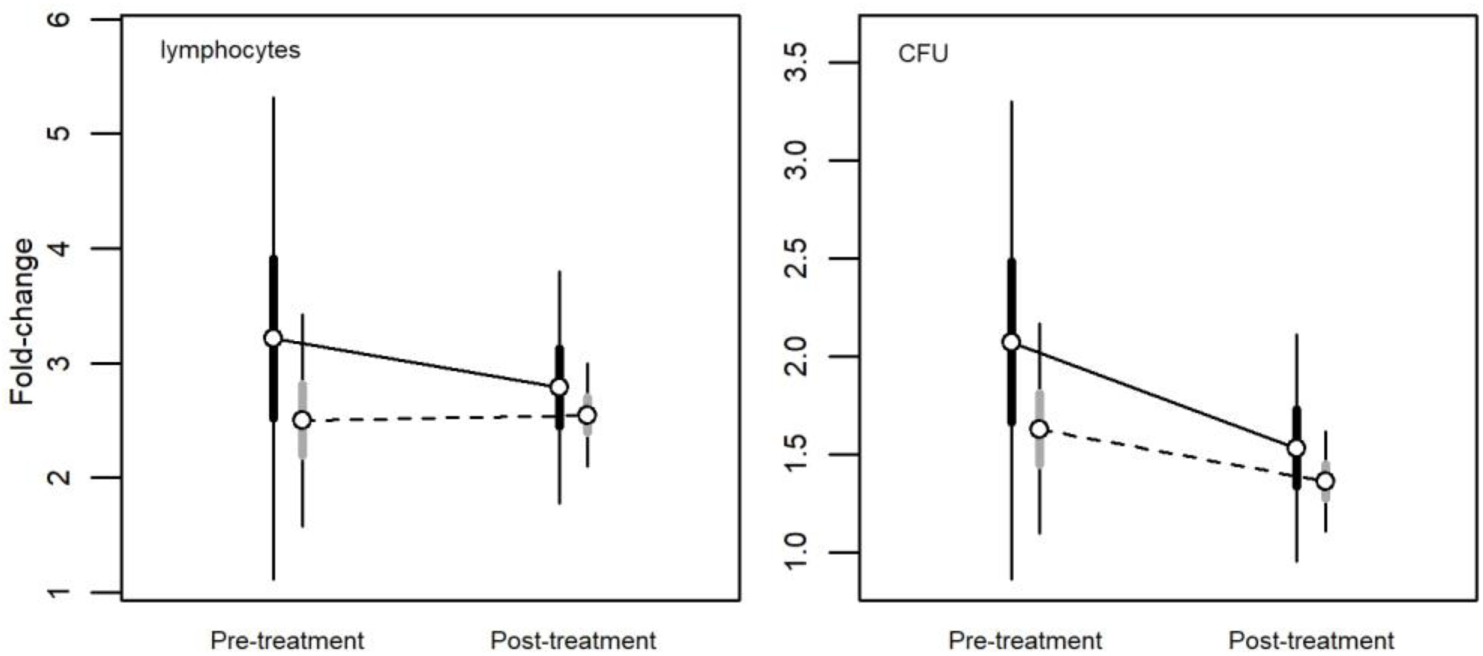
Bayesian model predictions for the immune-response of common wall lizards before (pre-treatment) and following (post-treatment) the testosterone supplementation according to morph (*black and gray bars* are for white and yellow morphs, respectively). *Lymphocytes*: the number of proliferating lymphocytes after PHA inoculation; *CFU*: colony- forming units following PHA inoculation (see text for details). Circles indicate HSM, and thick and thin lines represent HDI50 and HDI95, respectively.

**Table 3.** Posterior distributions for the immunological response of common wall lizards before (pre-treatment) and following (post-treatment) the testosterone supplementation according to morph. Values are fold-changes with respect the control. HSM and HDI95 estimates (between brackets) are shown.

| Variable |  | Pre-treatment | Post-treatment | Difference (post-pre) |
| --- | --- | --- | --- | --- |
| Lymphocytes | White | 3.22 (1.48 – 4.97) | 2.79 (1.94 – 3.63) | -0.43 (-1.50 – 0.653)<br>$P_{d<0} = 0.749$ |
| | Yellow | 2.50 (1.74 – 3.27) | 2.55 (2.17 – 2.92) | +0.04 (-0.43 – 0.52)<br>$P_{d<0} = 0.437$ |
| | **Difference (yellow-white)** | -0.71 (-1.84 – 0.40)<br>$P_{ylw<whi} = 0.855$ | -0.24 (-0.78 – 0.31)<br>$P_{ylw<whi} = 0.769$ | |
| CFU | White | 2.07 (1.05 – 3.09) | 1.53 (1.05 – 2.02) | -0.54(-1.17 – 0.09)<br>$P_{d<0} = 0.922$ |
| | Yellow | 1.63 (1.18 – 2.08) | 1.36 (1.15 – 1.58) | -0.27 (-0.55 – 0.01)<br>$P_{d<0} = 0.945$ |
| | **Difference (yellow-white)** | -0.45 (-1.11 – 0.21)<br>$P_{ylw<whi} = 0.869$ | -0.17 (-0.49 – 0.14)<br>$P_{ylw<whi} = 0.818$ | |

The CFU followed the same pattern of response as observed for total lymphocytes (Tab. 3). White males had higher immune-response than yellow males in both measurements (Tab. 3), but the difference was less marked (about half) after testosterone administration (Tab. 3, Fig. 2). Differently from total lymphocyte count, both morphs showed immune-suppression in response to testosterone administration, as the CFU sensibly decreased in the second measurement (Tab. 3, Fig. 2). However, the immune-suppressive effect was about twice in white than in yellow males (Tab. 3).

### Behavioural response and immune-function

The behavioural response clearly correlated with the immune-function in a similar way in both morphs, with males with higher immune-response entering less frequently in the half of the enclosure with the mirror, but remaining there for longer times and biting the mirrored image more frequently (Fig. 3, 4). Accordingly, Time increased with increasing immune-function for both total lymphocytes (Tab. 4), and CFU (Tab. 5), while the opposite occurred for Nmirror, which in general decreased with increasing immune-function (Tab. 4, 5). Further, males with higher immune-function bit more frequently the mirrored image (Tab. 4, 5), and the effect was particularly evident for the CFU in the post-treatment (Tab. 5).

**Figure 3.**
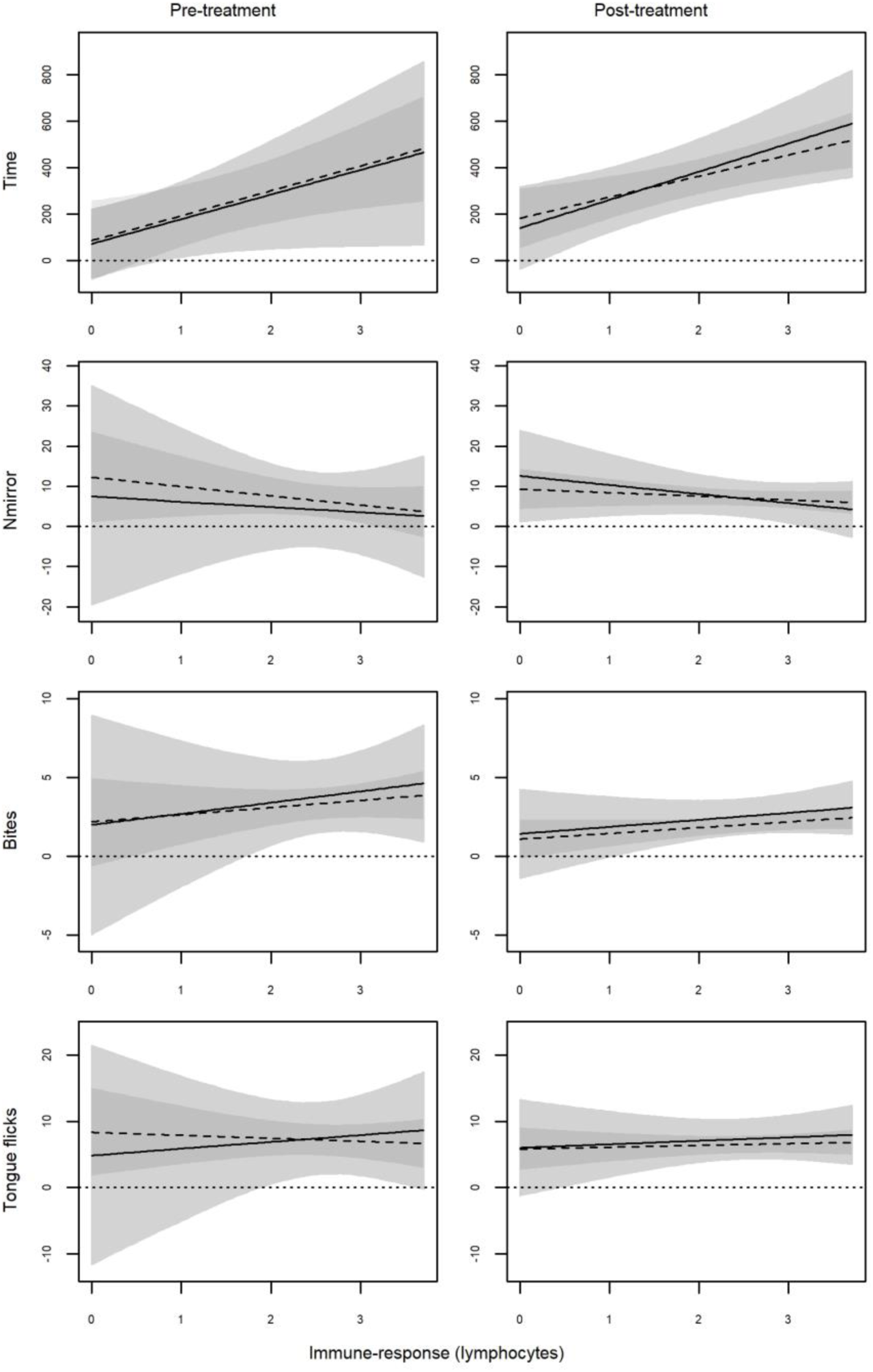
Bayesian model predictions for the relationship between behavioural response and immune-response (total lymphocyte count) of common wall lizards before (pre-treatment) and following (post-treatment) the testosterone supplementation according to morph (solid and light grey are for white morph and dashed and dark grey are for yellow morph). Lines indicate HSM, and areas represent HDI95.

**Figure 4.**
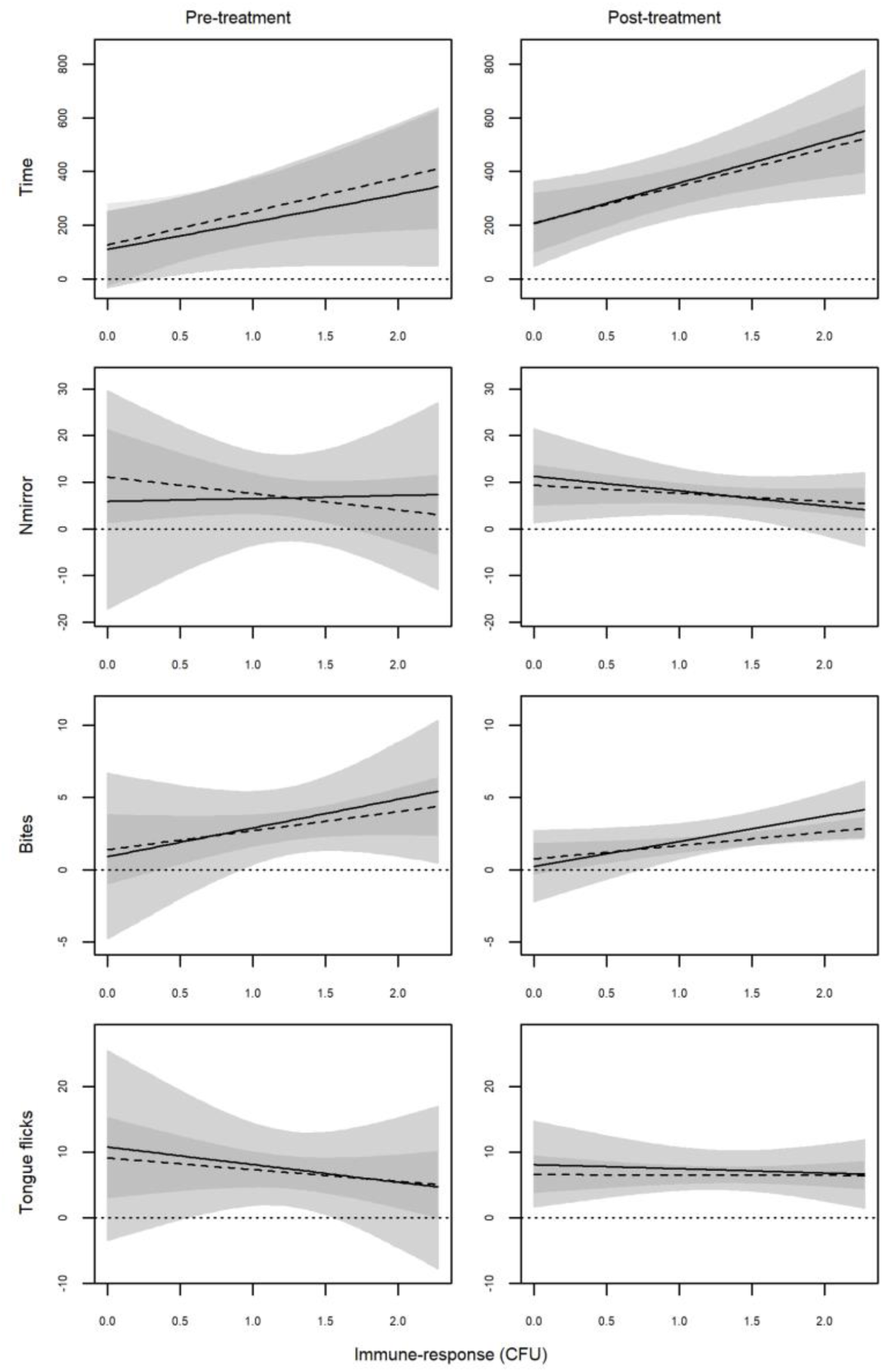
Bayesian model predictions for the relationship between behavioural response and immune-response (colony- forming units, CFU) of common wall lizards before (pre-treatment) and following (post-treatment) the testosterone supplementation according to morph (solid and light grey are for white morph and dashed and dark grey are for yellow morph). Lines indicate HSM, and areas represent HDI95.

**Table 4.** Posterior distributions for the slopes of the relationship between behavioural responses and immune-function as estimated by the total lymphocytes in the two morphs of the common wall lizard before (pre-treatment) and following (post- treatment) the testosterone supplementation. HSM and HDI95 (between brackets) estimates are shown.

| Variable |  | Pre-treatment | Post-treatment | Difference (post-pre) |
| --- | --- | --- | --- | --- |
| Time | White | 105 (-3 – 213)<br>$P_{\beta>0} = 0.945$ | 121 (39 – 203)<br>$P_{\beta>0} = 0.991$ | +16 (-61 – 93)<br>$P_{d>0} = 0.628$ |
| | Yellow | 106 (25 – 187)<br>$P_{\beta>0} = 0.984$ | 90 (38 – 143)<br>$P_{\beta>0} = 0.998$ | -16 (-75 – 43)<br>$P_{d<0} = 0.674$ |
| | **Difference (yellow-white)** | +1 (-80 – 83)<br>$P_{ylw>whi} = 0.515$ | -30 (-91 – 31)<br>$P_{ylw<whi} = 0.792$ | |
| Nmirror | White | -1.19 (-11.3 – 8.77)<br>$P_{\beta<0} = 0.547$ | -2.20 (-6.40 – 1.95)<br>$P_{\beta<0} = 0.816$ | -1.0 (-8.18 – 6.23)<br>$P_{d<0} = 0.589$ |
| | Yellow | -2.27 (-6.51 – 1.95)<br>$P_{\beta<0} = 0.806$ | -0.89 (-2.70 – 0.92)<br>$P_{\beta<0} = 0.789$ | 1.39 (-1.41 – 4.19)<br>$P_{d>0} = 0.795$ |
| | **Difference (yellow-white)** | -1.10 (-8.18 – 6.03)<br>$P_{ylw<whi} = 0.599$ | 1.32 (-1.42 – 4.07)<br>$P_{ylw>whi} = 0.782$ | |
| Bites | White | 0.70 (-1.82 – 3.25)<br>$P_{\beta>0} = 0.677$ | 0.44 (-0.58 – 1.47)<br>$P_{\beta>0} = 0.761$ | -0.26 (-2.12 – 1.57)<br>$P_{d<0} = 0.592$ |
| | Yellow | 0.44 (-0.57 – 1.47)<br>$P_{\beta>0} = 0.767$ | 0.36 (-0.08 – 0.81)<br>$P_{\beta>0} = 0.909$ | -0.08 (-0.77 – 1.55)<br>$P_{d<0} = 0.582$ |
| | **Difference (yellow-white)** | -0.26 (-2.09 – 1.55)<br>$P_{ylw<whi} = 0.589$ | -0.08 (-0.76 – 0.61)<br>$P_{ylw<whi} = 0.579$ | |
| TF | White | 1.06 (-5.07 – 7.02)<br>$P_{\beta>0} = 0.617$ | 0.56 (-2.15 – 3.12)<br>$P_{\beta>0} = 0.639$ | -0.52 (-4.82 – 3.85)<br>$P_{d<0} = 0.580$ |
| | Yellow | -0.46 (-2.92 – 1.94)<br>$P_{\beta<0} = 0.621$ | 0.29 (5.27 – 7.84)<br>$P_{\beta>0} = 0.660$ | 0.74 (-0.87 – 2.32)<br>$P_{d>0} = 0.778$ |
| | **Difference (yellow-white)** | -1.54 (-5.79 – 2.89)<br>$P_{ylw<whi} = 0.723$ | -0.28 (-1.95 – 1.48)<br>$P_{ylw<whi} = 0.605$ | |

**Table 5.** Posterior distributions for the slopes of the relationship between behavioural responses and immune-function as estimated by the CFU in the two morphs of the common wall lizard before (pre-treatment) and following (post-treatment) the testosterone supplementation. HSM and HDI95 estimates (between brackets) are shown.

| Variable |  | Pre-treatment | Post-treatment | Difference (post-pre) |
| --- | --- | --- | --- | --- |
| Time | White | 102 (-25 – 228)<br>**P<sub>β>0</sub> = 0.906** | 151 (22 – 279)<br>**P<sub>β>0</sub> = 0.972** | 49 (-63 – 159)<br>P<sub>d>0</sub> = 0.764 |
|  | Yellow | 124 (0.4 – 247)<br>**P<sub>β>0</sub> = 0.951** | 137 (54 – 221)<br>**P<sub>β>0</sub> = 0.996** | 13 (-86 – 114)<br>P<sub>d<0</sub> = 0.587 |
|  | **Difference (yellow-white)** | 22 (-84 – 126)<br>P<sub>ylw>whi</sub> = 0.638 | -14 (-108 – 83)<br>P<sub>ylw<whi</sub> = 0.582 |  |
| Nmirror | White | 0.50 (-17.0 – 17.5)<br>P<sub>β>0</sub> = 0.519 | -3.19 (-10.1 – 3.51)<br>P<sub>β<0</sub> = 0.781 | -3.68 (-16.2 – 9.11)<br>P<sub>d<0</sub> = 0.684 |
|  | Yellow | -3.67 (-11.1 – 3.63)<br>**P<sub>β<0</sub> = 0.795** | -1.73 (-4.68 – 1.17)<br>**P<sub>β<0</sub> = 0.837** | 1.95 (-3.34 – 7.33)<br>P<sub>d>0</sub> = 0.724 |
|  | **Difference (yellow-white)** | -4.19 (-15.8 – 7.70)<br>P<sub>ylw<whi</sub> = 0.721 | 1.45 (-3.03 – 6.01)<br>P<sub>ylw>whi</sub> = 0.701 |  |
| Bites | White | 1.94 (-2.26 – 6.29)<br>P<sub>β>0</sub> = 0.775 | 1.73 (0.06 – 3.40)<br>**P<sub>β>0</sub> = 0.955** | -0.22 (-3.43 – 2.88)<br>P<sub>d<0</sub> = 0.545 |
|  | Yellow | 1.30 (-0.47 – 3.08)<br>P<sub>β>0</sub> = 0.767 | 0.92 (0.20 – 1.64)<br>**P<sub>β>0</sub> = 0.983** | -0.37 (-1.67 – 0.90)<br>P<sub>d<0</sub> = 0.684 |
|  | **Difference (yellow-white)** | -0.64 (-3.67 – 2.31)<br>P<sub>ylw<whi</sub> = 0.684 | -0.80 (-1.91 – 0.31)<br>P<sub>ylw<whi</sub> = 0.882 |  |
| TF | White | -2.66 (-13.4 – 7.88)<br>P<sub>β<0</sub> = 0.664 | -0.57 (-5.06 – 3.81)<br>P<sub>β<0</sub> = 0.584 | 2.16 (-5.78 – 9.98)<br>P<sub>d>0</sub> = 0.672 |
|  | Yellow | -1.76 (-6.27 – 2.60)<br>P<sub>β<0</sub> = 0.746 | -0.02 (-1.94 – 1.82)<br>P<sub>β<0</sub> = 0.508 | 1.74 (-1.43 – 4.92)<br>P<sub>d>0</sub> = 0.818 |
|  | **Difference (yellow-white)** | 0.91 (-6.48 – 8.40)<br>P<sub>ylw>whi</sub> = 0.581 | 0.53 (-2.39 – 3.54)<br>P<sub>ylw>whi</sub> = 0.616 |  |

Morph specific effects of the testosterone administration on the relationship between aggressive behaviour and immune-response were detected for both Time and Nmirror. Indeed, the posterior probability of the three-way interaction term of the models deviated from zero in both total lymphocytes and CFU for Time (Pβ<0= 0.814 and Pβ<0= 0.741 respectively) and Nmirror (Pβ>0= 0.763; Pβ>0= 0.848 respectively). In the pre-treatment assessment, the relationship between the two behavioural responses and the immune-function was more stringent (i.e., narrower CI95%) in the yellow morph, while the white morph had a greater variability (Fig. 3, 4). After the testosterone administration, the variability in the white morph sensibly reduced, while not relevant changes occurred in the yellow one. Consequently, the dependence of the behavioural response on the condition of the immune system became equally stringent in the two morphs. We did not found any support for an equivalent three-way interaction for Bites with both lymphocytes (Pβ>0= 0.578) and CFU (Pβ>0= 0.546). However, we detected an appreciable positive main effect of the immune function (lymphocytes: Pβ>0= 0.677; CFU: Pβ>0= 0.775), a negative main effect of the treatment and a two-way interaction morph×immune-function, the last two only for the CFU (Pβ<0= 0.614, and Pβ<0= 0.641, respectively). In summary, the slope of the relationship between Bites and the immune-response is steeper in white with respect to yellow males, irrespective of the hormonal treatment (Fig. 3, 4). However, as for Time and Nmirror, the Bites vs immune-function relationship had a wider variance in white males than yellow one, and this difference no longer persisted after the hormone administration (Fig. 3, 4).

Finally, the effects of the immune-response on TF were contrasting and not fully coherent, if not opposite, when considering lymphocytes and CFU as immune-response measure (Fig. 3, 4). Indeed, TF increased with increasing total lymphocytes (Tab. 3), but, conversely, decreased with increasing CFU (Tab. 4). Morph specific effect appeared also for TF, but limited to lymphocytes (Pβ>0= 0.736; CFU: Pβ>0= 0.544). The general effect of the hormone administration on the TF vs immune-response relationship has been to reduce the difference in slope between morphs (Fig. 3).

## DISCUSSION

In this paper we manipulated plasma testosterone level (proven by TMs measured in droppings) to test if the white and yellow morphs of the common wall lizards represent two alternative and opposite solutions to the trade-off between immunity and the expression of secondary sexual characters depicted by the ICCH (Folstad and Karter 1992). Previous correlative results (Sacchi, Rubolini, et al. 2007; Sacchi, Scali, et al. 2017; Sacchi, Mangiacotti, et al. 2017; Coladonato et al. 2020) led us to hypothesise that white morph should follow an healthy strategy allocating more resources to the immune system at the expense of aggression in the defence of the territory. Conversely, we expected yellow males to follow a risky strategy, by allocating more resources in aggressive behaviour aimed at territory defence at the expense of effectiveness in immune response. We obtained a substantial, robust, experimental support for it.

First, before the hormone administration yellow males were immunosuppressed compared to the white ones, confirming morph specific immune-response previously observed in the species (Sacchi, Rubolini, et al. 2007; Sacchi, Mangiacotti, et al. 2017). According to the ICHH, increased plasma testosterone levels caused immunosuppression, but the effect was clearly morph specific, being sensibly more heavy in the white morph. Accordingly to the hypothesis of playing the healthy strategy, we showed white males not being able to cope with increased testosterone plasma levels. Their immune-system crashes when hormone levels are artificially raised beyond the maximum level allowed by the trade-off setting for maximum efficiency in immune function. Conversely, yellow males suffered a reduced immune-suppression as the trade-off setting for a greater expression of secondary sexual characters (i.e., the risky strategy) has predisposed the immune-system to better resist the immunosuppressive effects of increased plasma hormone levels. The administration of the hormone therefore has the overall effect of breaking out the correlation between the allocations into the immune function and secondary sexual character, with the final outcome of making the immune response in the two morphs similar.

Second, the administration of testosterone also depressed the intensity in the behavioural response in the post-treatment measurement, in a way that individuals become less aggressive than during the pre-treatment. Following the increase in plasma testosterone levels, males of both morphs bit less (i.e., reduced direct aggression), but stayed longer in the half enclosure with the mirror (i.e., increase indirect aggression). In other words, the hormone administration did not change the propensity of the individuals to face the intruder, but it rather caused a switch from the direct aggressive behavioural mode, based on fighting (riskier in terms of physical damage or injuries) to the indirect aggression behavioural mode, based on the threat, such as in a Mexican standoff (with a reduced risky of damage). The hormonal effect on the aggressive response was still more evident in the white morph, with the result that the differences between morphs disappeared following the hormonal treatment. Interestingly, the hormonal treatment did not have any relevant effects on the explorative behaviour (i.e., TF), supporting the strict relationship between aggressive behaviour and plasma testosterone levels.

Third, in both morphs the aggressive response, whether direct or indirect, strictly depended on the immune-function, irrespective of plasma testosterone levels. That is, only individual in better condition were able to sustain the costs of the territorial defence, in both morphs. This still reinforces the idea that immunity and aggression are linked in a trade-off and that the differences in immune response and aggressive behaviour that emerge between white and yellow morphs are due to the evolution of alternative responses to the impossibility of simultaneously maximizing the two functions.

We therefore supplied experimental evidence of the existence of two morph-specific strategies in common wall lizards depending on the investment in territorial aggression or in a longer survival. Nonetheless, this does not mean that yellow males are more aggressive than white ones in absolute terms, but in a relative sense. Indeed, we showed mean reaction norms for immune response, aggressive behaviour and plasma testosterone levels are similar between morphs, but it is the seasonal pattern that varies differently between morphs (Sacchi, Scali, et al. 2017; Sacchi, Mangiacotti, et al. 2017; Coladonato et al. 2020; Sacchi et al. 2021). Yellow males maintain higher T levels and display more aggressively over the beginning of the season, but display a stronger subsequent decline until having lower T levels and aggressive response than white males later in the season. Increased aggressive behaviour in the early part of the season means more clashes among individuals, at the cost of lower long-term survival due to both lower immune response and higher predatory risk (Marler and Moore 1988; Sacchi et al. 2009) to the benefits of white males who choose the conservative strategy. On the other hand, increased immune function means longer survival at the cost of a reduced competitive ability for obtaining the best territories, to the benefit of yellow males who may achieve more mating attempts. From an evolutionary point of view, the yellow strategy could be seen as a way to obtain a high reproductive success but over a short time interval (lifespan), while the white strategy should be a way of achieving less reproductive success but over a longer time interval. If the overall balance in terms of fitness between the two alternatives is equivalent over long time intervals, the selection should keep the strategies in balance over time, without one prevailing over the other. Hence, understanding how the two strategies link the reproductive opportunities, and therefore the possibility of mating with females, is crucial to figure out how they evolved and persisted within the same population. Females in Mediterranean lizards normally lay up two and often three annual clutches (the larger individuals), from early spring to late summer (*Podarcis ionica*, (Chondropoulos and Lykakis 1983); *P. atrata*, (Castilla and Bauwens 2000a); *P. lilfordi*, (Castilla and Bauwens 2000b); *P. bocagei*, (Galán 1997)), as well for common wall lizards (Barbault and Mou 1988). In Northern Italy the first clutch deposition of *P. muralis* occurs between late April and the beginning of May, while the second one occurs between late May and mid-July (Sacchi et al. 2012). In common wall lizards, as in other Mediterranean species, clutch size typically declines as the reproductive season progresses (Barbault and Mou 1988; Castilla and Bauwens 2000b). This seasonal variation is induced by differences in the proximate source of the energy allocated to different annual clutches. Furthermore, yolk production for the first clutch mainly derives from fat reserves stored before hibernation, whereas the energy shunted to subsequent clutches derives from recent food intake (Braña et al. 1992). This makes the first clutch a more remunerative and predictable resource for males in term of fitness compared with the second clutch, and even more the third one. Therefore, being very aggressive and defending the best territories in the first part of the season allows yellow males to access to the most profitable females, and therefore achieve the relative highest reproductive success within a single season. Nevertheless, the high breeding success pays the cost of a shorter life, which for *Podarcis* lizards means no more than two consecutive seasons (Barbault and Mou 1988; Altunışık et al. 2016). At the opposite, reducing the risk of damage or injury in the period of maximum competition and investing in longevity may actually force white males to focus on less profitable, and more unpredictable clutches having in return an additional time compared with yellow males. Hence, the variability of clutches laid by females along the season combined to the impossibility for males to compete all over the reproductive season might be the primary evolutionary driver promoting the setting of the trade-off inherent ICHH towards the dual solution of a risky vs conservative strategy.

Adopting a different colour badge depending on strategy can help male morphs to efficiently recognise the strategy of rivals and modulate their own behaviour, reducing the risk to income in unnecessary and potentially dangerous fighting. In fact, the scenario we have just depicted above allows us to predict that for a given male a rival adopting its own same strategy is potentially riskier, compared to another one who adopts a different strategy. Many species of vertebrates show a morph-specific aggressive response. For example, males of the cichlids fish direct more aggressive attempts towards similarly coloured opponents (Dijkstra et al. 2007) and similar outcomes have been reported for the polymorphic sparrow, *Zonotrichia albicollis* (Horton et al. 2012). Morph specific aggression also occurs in lizards as reported for the ornate tree lizard (*Urosaurus ornatus*), and the tawny dragon *Ctenophorus decresii* (Hover 1985; Yewers et al. 2016). Recently, we experimentally showed that aggression is more common during homomorphic than heteromorphic contests also in common wall lizards, and such a kind of interactions controls the patterns of spatial distribution of morphs (Scali et al. 2021). By manipulating testosterone plasma levels, we supplied experimental evidence in support to the hypothesis that colour morphs in this species have evolved as a status badge to inform conspecific about the strategies played by the signaller, and decrease unnecessary conflicts among different colour morphs.

